# A synthetic system for asymmetric cell division in *Escherichia coli*

**DOI:** 10.1101/436535

**Authors:** Sara Molinari, David L. Shis, Shyam P. Bhakta, James Chappell, Oleg A. Igoshin, Matthew R. Bennett

## Abstract

We describe a synthetic genetic circuit for controlling asymmetric cell division in *E. coli* in which a progenitor cell creates a differentiated daughter cell while retaining its original phenotype. Specifically, we engineered an inducible system that can bind and segregate plasmid DNA to a single position in the cell. Upon cell division, co-localized plasmids are kept by one and only one of the daughter cells. The other daughter cell receives no plasmid DNA and is hence irreversibly differentiated from its sibling. In this way, we achieved asymmetric cell division through asymmetric plasmid partitioning. We then used this system to achieve physical separation of genetically distinct cells by tying motility to differentiation. Finally, we characterized an orthogonal inducible circuit that enables the simultaneous asymmetric partitioning of two plasmid species, resulting in cells that have four distinct differentiated states. These results point the way towards engineering multicellular systems from prokaryotic hosts.

## Introduction

Synthetic biology enables fundamental studies of biology^1,2^ and the construction and characterization of genetic systems from the ground up^3–5^. Synthetic microbial organisms now hold promise for complex applications such as targeting tumors^6^, antibiotic and gene therapies^7,8^, microbiome manipulation^9^ and geoengineering^10^. However, synthetically engineered bacterial systems are relatively simple compared to complex multicellular organisms. While some synthetic bacteria can produce population-scale behaviors such as pattern formation^11–14^, robust synchronized oscillations^15–17^ and growth rate control^18^, no synthetic bacteria can compare to the highly coordinated activities of multicellular organisms.

One method found in nature for creating complex spatially distributed systems is cellular differentiation via the process of asymmetric cell division. Asymmetric cell division enables the development of different cell types throughout the organism to specialize by partitioning biochemical or physical tasks throughout the organism. In multicellular eukaryotes, cellular differentiation is generally achieved through complex regulatory networks that utilize transcriptional regulation, post-transcriptional modifications, and chromatin remodeling^19–21^. To differentiate asymmetrically, a progenitor cell (such as a stem cell) senses chemical cues in the environment to alter the transcriptional landscape in the daughter cell. This transcriptional rearrangement of the daughter cell is complete enough that de-differentiation is rare and reproducible in the lab only through the very specific and simultaneous induction of many genes^22^.

Several attempts have been made to create synthetic bacteria that have multiple stable transcriptional profiles^3,23^. The first of these was the corepressive toggle switch in *E. coli* designed by Gardner and colleagues^3^. However, the corepressive toggle does not create irreversibly differentiated cell types. It is sensitive to intrinsic and extrinsic sources of gene regulatory noise^3,24–27^, and can only transiently maintain one state before stochastically switching back to the other. Others have used recombinases^28^ to alter DNA sequences permanently, or controllably alter the copy numbers of plasmids to affect gene regulation and cell differentiation^29,30^.

Here, we repurposed two key elements of the chromosome partitioning system (*par*) of *Caulobacter crescentus*^31^ to create asymmetric cell division and irreversible differentiation in *E. coli*. The *par* system is common in prokaryotes and is principally responsible for partitioning low copy number plasmids or chromosomes upon cell division^32^. The *par* systems generally rely on the interaction of three elements: a centromere-like cis-acting sequence, *parS*, present on the plasmid/chromosome; a centromere-binding trans-acting protein, ParB; and an NTPase, usually called ParA^33^. The initial step is the formation of the partition complex, in which all the copies of the plasmid or chromosomal DNA are gathered together by the cooperative binding of many ParB proteins at the *parS* site. The ParB protein encoded by *C. crescentus* is very similar to the one present in the Type Ia *par* systems^32^ and contains a helix-turn-helix DNA binding motif that recognizes and cooperatively binds the cognate sequence, *parS*. A combination of homodimerization and nonspecific DNA-protein interactions leads to the formation of a nucleoprotein complex spreading for several kilobases around the *parS* site^34,35^. The exact mechanism of the ParB nucleation around *parS* is still largely unclear, but one model suggests a “nucleation and caging” mechanism in which a core of tightly bound ParB dimers forms around *parS*^36^.

In this study, we show that by repurposing the *par* system of *C. crescentus* in *E. coli* (which does not contain a genomic *par* system^37^) one can control the asymmetric partitioning of plasmids. We then used this system to achieve physical separation of genetically distinct cells by tying motility to differentiation. We further show that repurposing the *par* system from the F plasmid facilitates an orthogonal pathway for inducible asymmetric plasmid partitioning (APP). Using these two pathways, we engineered a genetic circuit in which two different plasmids can be independently partitioned to create four distinct cell types.

## Results

### APP in *E. coli*

The circuit we constructed for APP in *E. coli* is illustrated in Fig. 1a. It consists of two elements of the *par* operon from *C. crescentus:* the centromere-like site *parS* and the centromere-binding protein encoded by *parB*^33^. We cloned the cis-acting *parS* sequence onto a plasmid that we refer to as the “target plasmid.” On a second plasmid, which we call the “regulatory plasmid” we cloned the gene encoding ParB fused to super folder yellow fluorescent protein (sfYFP) and a hydrophobic leader peptide (MKAIFVLKHLNHAKETS). This gene (*lp-sfyfp-parB*) was placed under the control of an arabinose-inducible promoter. When present, ParB binds to and forms a nucleoprotein complex around the *parS* site^31^. This characteristic of ParB consolidates copies of the target plasmid into a single cluster as shown in Figs. 1b and 1c. Upon induction of the system, one daughter cell ultimately inherits the nucleoprotein oligomer, facilitating the asymmetric partitioning of the target plasmid. The other septation partner loses the target plasmid and becomes terminally differentiated. In this way, asymmetric cell division happens through APP (Fig. 1d).

**Figure 1.**
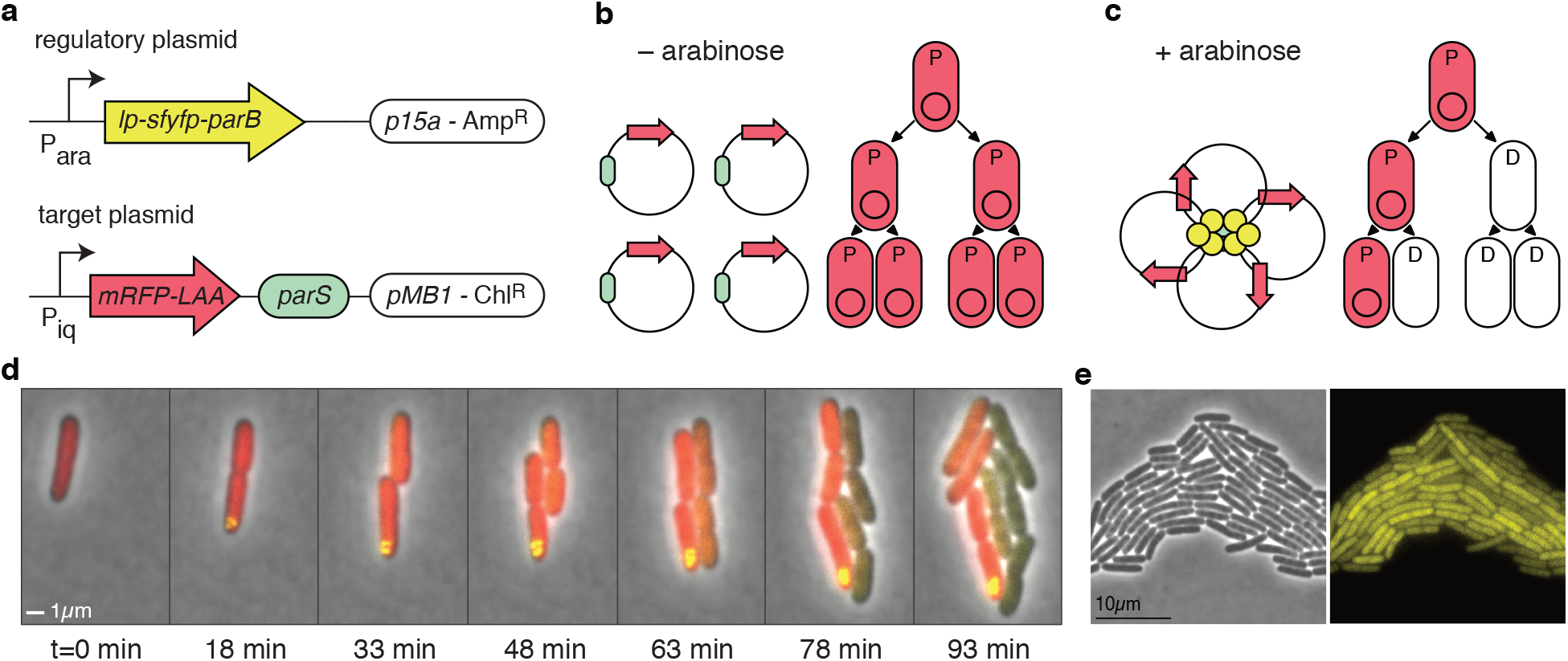
Asymmetric plasmid partitioning in *E. coli*. **(a)** The APP network consists of two plasmids: the regulatory plasmid containing *lp-sfyfp-parB* (lp: leader peptide) under the control of an arabinose inducible promoter and the target plasmid containing the *parS* sequence and a red fluorescent protein gene. **(b)** In the absence of arabinose, there is no expression of *lp-sfyfp-parB*, so target plasmids are free to diffuse in the cell. This means that they segregate roughly symmetrically in the population in which all cells are progenitor cells (denoted by “P”). **(c)** When arabinose is present, lp-sfYFP-ParB binds *parS* on the target plasmid, forming a nucleoprotein complex that gathers all copies of the plasmid together. In this case, cells begin to asymmetrically divide, giving rise to differentiated cells (denoted by “D”). **(d)** Time-lapse fluorescence microscopy of cells undergoing asymmetric plasmid partitioning. Shown at time t=0 is a single progenitor cell that is in the presence of arabinose. A nucleoprotein complex quickly forms (yellow punctum), and subsequent daughter cells lose the target plasmid. The inherited red fluorescent protein in the daughter cells quickly decays through dilution and proteolysis. **(e)** Phase contrast (left) and yellow fluorescence microscope images of cells containing only the regulatory plasmid, encoding *lp-sfyfp-parB*, induced with 0.2% arabinose.

To characterize the induction of APP, we first utilized single cell fluorescence microscopy by tracking the nucleoprotein complex around the *parS* sequence, which appears as a yellow fluorescent punctum. To track the segregation of target plasmid DNA, we cloned a gene encoding a red fluorescent protein (*mRFP*) with a C-terminal degradation tag (AANDENYALAA)^38^ onto the target plasmid. We next followed the proliferation of red fluorescence in dividing cells using fluorescence microscopy. Cells transformed with both the regulatory plasmid and the target plasmid were cultured on an agarose pad perfused with arabinose and imaged over time (see Methods). At the beginning of cell growth, several small florescent foci formed inside the cells at apparently random positions; these foci then segregated randomly to one or both daughter cells (Fig. 1d and Supplementary Fig. 1). In the latter case, this correlates with the observation that both daughter cells retain red fluorescence, suggesting inheritance of target plasmid DNA. However, as the puncta of YFP fluorescence become larger and consolidate, segregation of the nucleoprotein complex to only one daughter cell upon cell division becomes the norm. In this case, red fluorescence tightly correlates with the presence of a single punctum of YFP fluorescence. The presence of fluorescent puncta suggests the presence of the target plasmid inside the cell (Fig. 1d, e). The loss of fluorescent puncta, hence, suggests that the cell has lost target plasmid DNA. In agreement with this hypothesis, single cell microscopy shows that daughter cells that did not inherit fluorescent puncta (and presumably the plasmid-ParB complex) also rapidly degrade the red fluorescence signal. Furthermore, once red fluorescence was lost, we never observed it to be recovered. This contrasts with the maintenance of red fluorescence in cells that retain YFP puncta. A representative example of this process is shown in Fig. 1d.

The nucleoprotein complex that forms around *parS* may have a silencing effect on neighboring genes^39^. To test whether this occurs in our system, we cloned a target plasmid in which *parS* is only 56 bp from the promoter driving *mRFP* (the target plasmid we used to acquire the pictures in Fig. 1d has a spacer about 3.2 kb long that separates the *parS* site from the *mRFP* promoter). We measured the red fluorescence of 12 mother cells every 9 min and confirm the silencing effect on genes in close proximity to *parS* (Supplementary Fig. 2a). The absence of the spacer does not affect the efficiency of APP (Supplementary Fig. 2b).

In addition to single cell fluorescence microscopy, we also tracked the proliferation of target plasmid DNA in populations by first growing them in flask (with or without inducer) and then plating them onto LB agar plates (with or without chloramphenicol, the selective antibiotic for target plasmid), as depicted in Fig. 2a (see Methods). In the absence of arabinose, APP does not occur, so target plasmid DNA segregates normally. Hence, chloramphenicol resistance is present in every cell, and the number of colony-forming units (CFUs) at each stage of growth does not depend on the presence or absence of chloramphenicol in the plate (Fig. 2b, black and blue curves). The results were very different, however, when arabinose was included in the liquid culture. In that case, CFU counts were near the uninduced counts when plated onto plates lacking chloramphenicol. This is expected, as both cells with (progenitor cells) and without (daughter cells) the target plasmid should grow normally (Fig. 2b, green curve). When plated onto selective plates, however, CFU counts were drastically lower (Fig. 2b, red curve).

**Figure 2.**
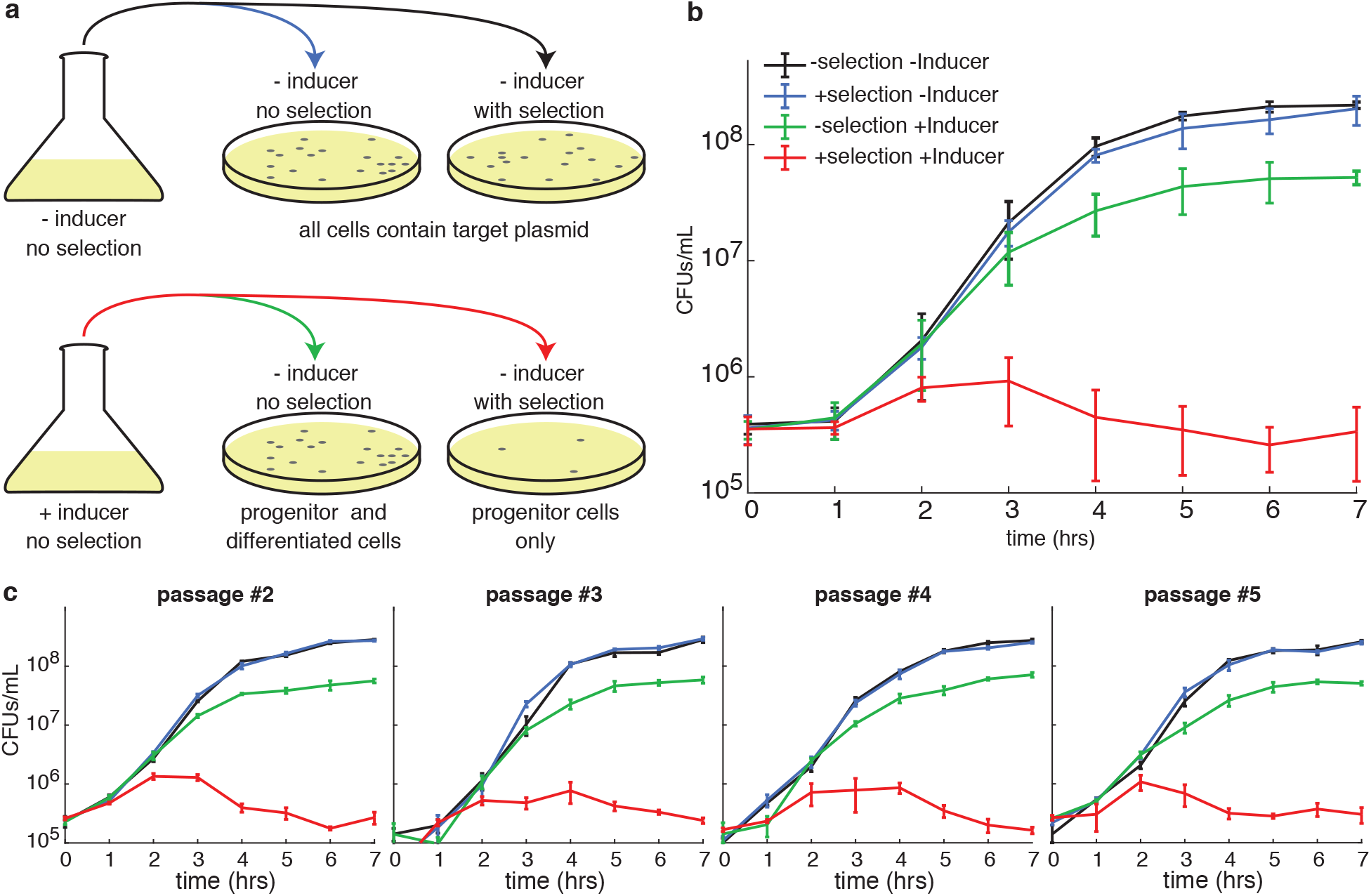
Progenitor cells can undergo multiple rounds of APP. **(a)** Pictorial description of the plating assay. Liquid cultures without (top) and with (bottom) inducer were plated onto agarose with and without selection for the target plasmid. Without inducer (top) the number of CFUs should be the same independent of the antibiotic. With inducer, however, only progenitor cells will grow on the plate that includes antibiotic. **(b)** CFUs as a function of time for each of the four conditions described in (a). In the case of the uninduced culture (black and blue lines) there is almost no difference between cells plated with or without chloramphenicol. For the induced culture (green and red lines), however, only the cells plated without chloramphenicol show normal growth (green line), whereas the number of cells able to grow on chloramphenicol (progenitor cells) remain roughly constant over time. Error bars represent standard deviation of three biological replicates measured in technical triplicate. **(c)** CFUs as a function of time for four sequential re-inoculations of cells from (b). Error bars represent standard deviation of a single biological replicate measured in technical triplicate.

We observed a reduction in overall CFU counts when the APP network is induced at 0.2% arabinose, suggesting a non-negligible fitness cost associated with induced *lp-cfp-parB* expression (Fig. 2b). Note that for all the experiment not involving single cell microscopy, we switched to *lp-cfp-parB* to make sfYFP available for later microscopy experiments, however, both *lp-cfp-parB* and *lp-sfyfp-parB* behaved similarly in our assays (see below).

We next tested the robustness of our system by performing multiple cycles of induced APP in sequence. To do this, we picked and regrew a colony from a 7-hour plate with chloramphenicol with cells from the induced culture (the rightmost point of the red curve in Fig. 2b). On this plate all cells should be progenitor cells and have both plasmids of the APP system. On the following day, we repeated the above process of induction of APP and plating. CFU counts for each case were similar to those obtained with the first induction, as were those on subsequent repetitions of the experiment (Fig. 2c). This means that even after a round of APP, the progenitor cells were able to recover and undergo subsequent rounds of APP.

As a further confirmation of the dynamics of the target plasmid within the induced and uninduced populations, we also analyzed the plasmid content of both using qPCR to examine the dynamics of APP in the growing populations. As shown in Supplementary Fig. 3, the ratio of target plasmid (segregating asymmetrically) to the regulatory plasmid (segregating symmetrically) decreases over time in the induced population. In contrast, the uninduced population shows a roughly constant ratio of the two plasmids over time.

To verify that the target plasmid is lost in differentiated cells, we picked 12 colonies from the induced population plated without chloramphenicol (where the majority of cells are expected to have lost the target plasmid) and measured the amount of target plasmid DNA using qPCR. They were all positive for the presence of the regulatory plasmid, but none show any amplification of the target plasmid for the first 30 cycles of amplification in the qPCR (Supplementary Table 1a). Moreover, no colony grew on chloramphenicol-containing LB but all of them grew on plates lacking chloramphenicol. To verify that the loss of target plasmid is solely due to the accumulation of ParB protein, we tested the efficiency of APP in a system containing either a target plasmid with a deleted *parS* domain or a regulatory plasmid missing ParB (expressing only *lp-cfp*). We observed a negligible loss of target plasmid in both cases (Supplementary Fig. 4).

From the data presented in Figs. 2b and 2c we noticed a certain degree of toxicity in the expression of the ParB construct, in both progenitor and differentiated cells. We tested whether a longer exposure to this protein increases the toxicity effect. Supplementary Fig. 5 shows that 23 h after induction (24 h after inoculation) the culture has a nearly identical ratio and absolute number of progenitor and differentiated cells. This also shows that the complex is stable long after cells entered stationary phase (5–7 h after inoculation) and stop producing ParB.

We next investigated the role of the different domains of the *lp-(cfp/sfyfp)-parB* genes in facilitating APP. We hypothesized that ParB alone, although capable of forming a nucleoprotein complex around *parS*^35^, could not build a sufficiently strong protein-DNA cluster capable of gathering all copies of the plasmid together. Both the leader peptide and the fluorescent protein may help stabilize ParB-ParB interactions. The leader peptide at the N-terminus of the construct is 17 amino acids long, the first 7 of which form a hydrophobic block with a grand average of hydropathicity (GRAVY) of 2.16^40^, and it is known that some fluorescent proteins have a natural tendency to oligomerize^41^.

We tested various leader peptides and fluorescent tags with different properties and compared the resulting efficiency to the *lp-cfp-parB* construct, which is the one used to generate the data in Figs. 2b,c. Supplementary Fig. 6 shows the fraction of progenitor cells in induced and uninduced cultures. A lower fraction of progenitor cells in the population is indicative of a more efficient APP. The deletion of either the leader peptide or *cfp* from the original construct results in a very mild activation of APP, whereas the expression of *parB* alone shows no APP at all. If we substitute the leader peptide with a sequence with different properties, we can change the efficiency of the system: a glycine-serine peptide (GGGGS)_4_ which has a GRAVY index of −0.48 and no polarity, fused N-terminally to *cfp-parB* is not able to facilitate APP. In contrast, fusing a GCN4 leucine zipper mutant (PDB 1CE9), known for its dimerization properties^42^, to either *cfp-parB* or *parB* allowed for an efficiency of APP that is comparable to the original. In addition, by varying the fluorescent protein we obtained a broad range of APP activation. As expected, substituting CFP with sfYFP, due to their similar ability of oligomerizing, does not result in a significant change in efficiency compared to the CFP system. Replacing CFP with mCherry, a highly monomeric fluorescent protein^41^, still allows a small amount APP, presumably because mCherry does not provide the extra oligomerizing effect that CFP does. However, a construct containing a strong oligomerizing protein, DsRed, shows a lower fraction of progenitor cells than the original system in the induced population^41^.

### Tuning the efficiency of APP

We next explored potential strategies for tuning ligand inducible APP. To do this, we tested two other versions of the target plasmid that contain different origins of replication with different copy numbers (pUC and pSC101) in addition to the original version containing pMB1. The pUC origin of replication is a mutant pMB1 that confers a much higher copy number (~300–500 copies per cell^43^) compared to wild type (~10–20 copies per cell^44^). We specifically wanted to know if a target plasmid with a high copy number would aggregate and segregate as efficiently as the one with the pMB1 origin. The pSC101 origin confers a low copy number (~5 copies per cell) and is actively partitioned by ParA of *E. coli’s* SMC complex^45^. We wanted to know if the active segregation mechanism of pSC101 would interfere with the APP systems ability to aggregate the target plasmid. All three versions of the target plasmids were then tested with various amounts of inducer. As shown in Fig. 3, the fraction of progenitor cells (as measured by the plating assay) decreases with increasing inducer concentrations for each type of plasmid. This is true even in the case of pSC101, though there is a considerable amount of APP activation even when the system is uninduced.

**Figure 3.**
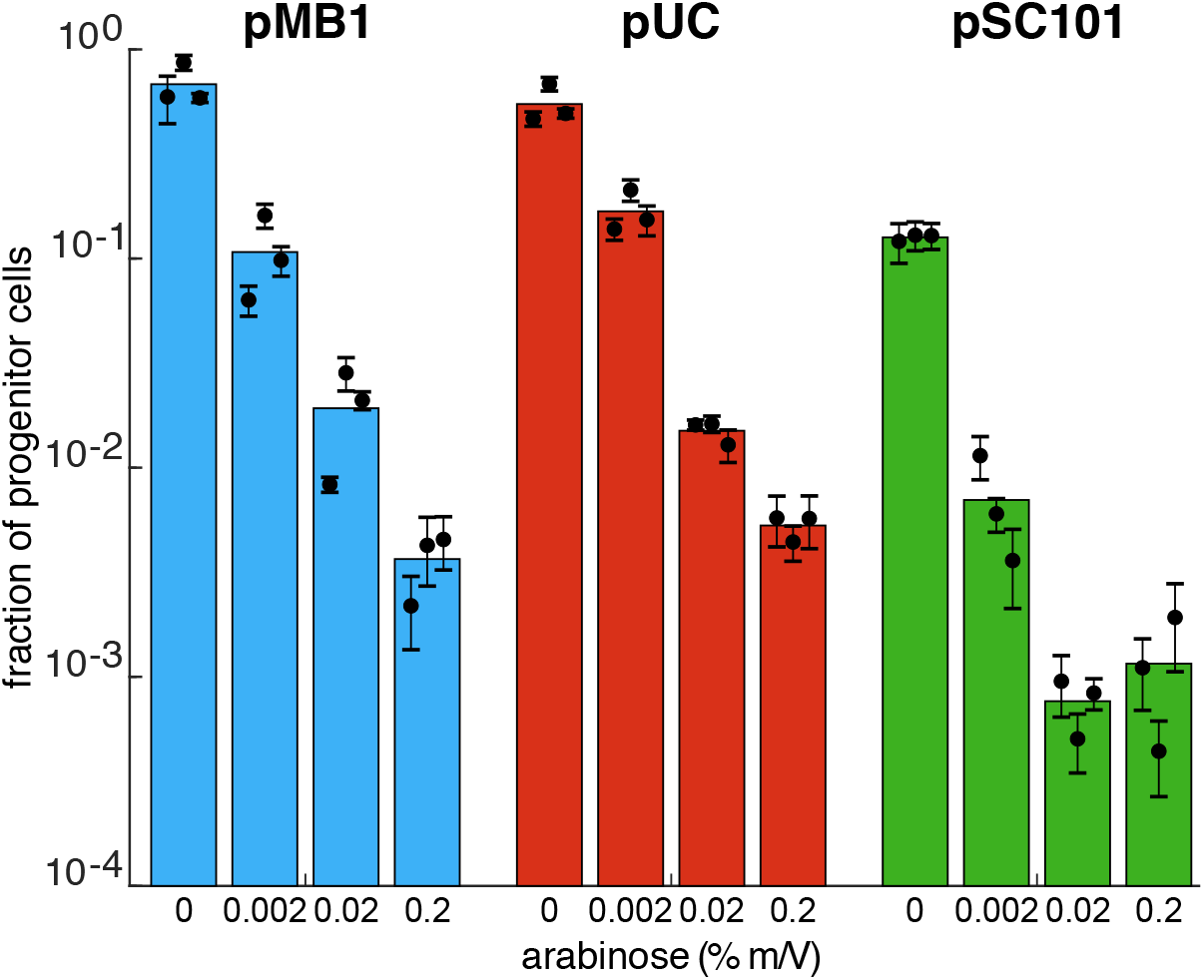
APP with different origins of replication. Fraction of progenitor cells as a function of inducer concentration using a target plasmid with different origins of replication as measured by plating assay: pMB1 (blue) has a medium copy number (~15-20) and a passive mechanism of partitioning, pUC (purple) has a high copy number (~300-500) and passive mechanism of partitioning, pSC101 (green) has a low copy number (~5) and an active mechanism of partitioning. Results show that the efficiency of APP can be modulated by varying the amount of inducer. Colored bars represent the average of all trials, black dots represent the average of each biological replicate and their error bars the standard deviation of technical triplicates.

### Physical separation of differentiated cells

We next demonstrated that it is possible to use APP to physically separate progenitor from differentiated cells by linking motility to the presence/absence of the target plasmid. To do this, we used a motile strain of *E. coli* (HCB84) carrying a *motA* mutation known to disrupt motility^46^. The motility phenotype can be restored by overexpressing *motA* on a plasmid. To link motility to APP, we created the plasmids shown in Fig. 4a. Essentially, the target plasmid now contains a repressor, PhlF, that downregulates *sfyfp* and *motA* encoded on a third plasmid. When APP begins (via induction of *lp-cfp-parB* by IPTG), the target plasmid is lost in daughter cells, PhlF levels decrease, and *motA* and *sfyfp* become derepressed – allowing motility and fluorescently labeling differentiated cells.

**Figure 4.**
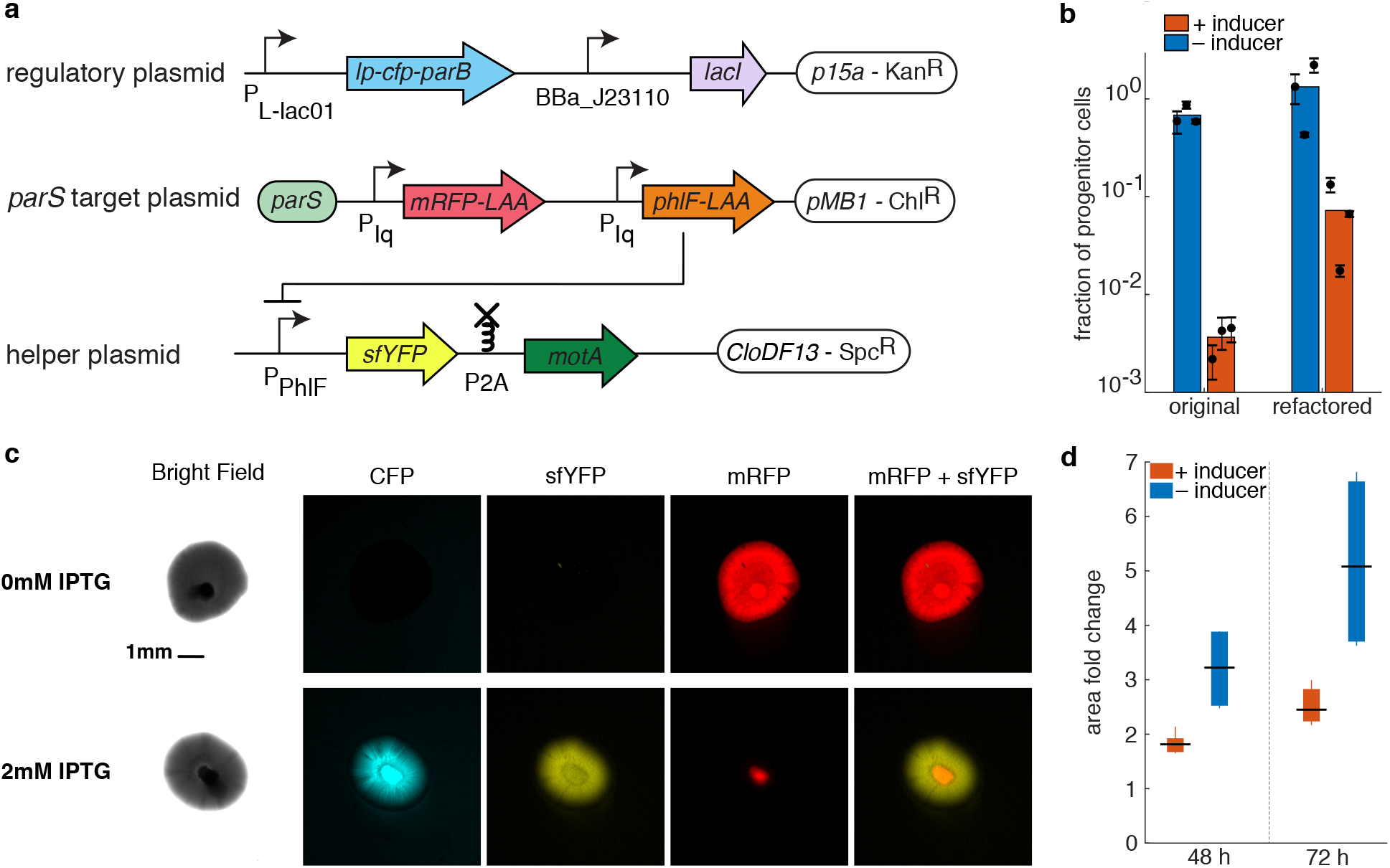
Physical separation of genetically different cells. **(a)** APP circuits refactored to be functional in HCB84 *E. coli* strain. It consists of three plasmids, out of which only one is the target plasmid and will, therefore, asymmetrically partition. **(b)** The graph shows the efficiency of the refactored circuit in *E. coli* HCB84 compared to the original system in JS006 ALT as measured by plating assay. Colored bars represent the average of all trials, black dots represent the average of each biological replicate and their error bars the standard deviation of technical triplicates. **(c)** Pictures of colonies growing in semi-solid agarose plates (0.3%) with (bottom panel) and without (upper panel) inducer. Columns show different fluorescence channels of the same image. Scale bar is shown only at time on the bright field picture of the uninduced condition and is the same for every image. **(d)** Area fold change of colonies grown on agarose with and without inducer. The area of each colony at time 48 and 72 h was normalized to the area of the same colony 24 h after the inoculation. Data shown as standard box plot.

Fig. 4b shows that the fraction of progenitor cells in induced culture expressing the new circuit (expressed in *E. coli* HCB84) are considerably higher compared to the original system in JS006 ALT. Nevertheless, the number of differentiated cells is about 15-fold higher than the progenitors. This ratio is enough to clearly observe the physical separation of genetically differentiated cells on semi-solid agarose. We inoculated saturated cultures of cells transformed with the APP circuit (Fig. 4a) into semi-solid agarose (see Methods). Figure 4c shows the growth of a single colony 48 hours after inoculation into agarose without (right panel) or with IPTG (left panel). When cells were exposed to IPTG they expressed the ParB construct (as indicated by cyan fluorescence) and initiated the production of differentiated cells (identifiable by yellow fluorescence), which became motile and migrated toward the outer edge of the colony.

In contrast, colonies grown in the absence of IPTG did not produce differentiated cells, and the colony appears to express only *mRFP*, indicative of it consisting of progenitor cells only. More examples of colonies imaged in the exact same conditions, in both inducing and noninducing media, are shown in Supplementary Fig. 7.

We measured the area of six induced and six uninduced colonies 24, 48 and 72 h after their inoculation into soft agarose. Areas at time 48 and 72 h were normalized against the first time point (24 h). In this way, we compared the area fold change in induced and uninduced colonies. As shown in Fig. 4d, induced colonies grew twice more than uninduced colonies. This observation is in agreement with the fact that induced colonies produce differentiated cells that are motile, while uninduced colonies are only made of non-motile progenitor cells.

### An orthogonal system for multiple differentiated states

Lastly, we explored the possibility of expanding the potential of APP by repurposing a second orthogonal APP circuit into *E. coli*. Our end goal was to build a circuit capable of independently partitioning two different plasmids upon the induction of two separate trans-acting proteins. In this way, one could differentiate an initial isogenic strain into four different cell types. To do this, we first replaced *parB* on the regulatory plasmid with *sopB*, and the *parS* site on the target plasmid with *sopC*. These two elements are from the F plasmid, and have native functions similar to their counterparts^47^.

Just as with the ParB/*parS* system, the fraction of progenitor cells decreases in the SopB/*sopC* system as a function of increasing amount of inducer (Supplementary Fig. 8a), and subsequent rounds of APP were possible, provided the inducer concentration was not too high (Supplementary Fig. 8b). We next assessed the orthogonality of the two networks by comparing the results all the four possible combinations of the two plasmid-gathering proteins and two cis-acting sequences (Supplementary Fig. 8c). Only the correct pairing of the APP network elements resulted in APP, whereas the mispaired combinations ParB/*sopC* and SopB/*parS* did not. We also observed that the construct lp-SopB is able to facilitate APP even in the absence of the fluorescent tag, however, the efficiency of APP decreases significantly compared to the original SopB construct, as shown in Supplementary Fig. 8d (the construct still has the leader peptide). This is in agreement with our hypothesis on how each piece of the leader peptide–fluorescent tag–centromere binding protein construct affect the APP efficiency.

To construct the two-plasmid APP circuit, we combined the two synthetic APP pathways together by repurposing the four genetic elements into a new circuit made of three plasmids: two target plasmids (each containing one of the two centromere-like sequences, *parS* and *sopC*) with chloramphenicol and spectinomycin resistance, and a regulatory plasmid with *parB* and *sopB* being driven by arabinose and IPTG inducible promoters, respectively (Fig. 5a). With this new circuit, the progenitor cells can differentiate in several ways (Fig. 5b). If either inducer is used alone, progenitor cells should begin to produce one of two partially differentiated cell types that lack one of the target plasmids. If both inducers are used simultaneously, progenitor cells produce terminally differentiated cells lacking both target plasmids. Finally, if one sequentially induces the system with first one inducer and then the other, partially differentiated cells should begin to produce terminally differentiated cells.

**Figure 5.**
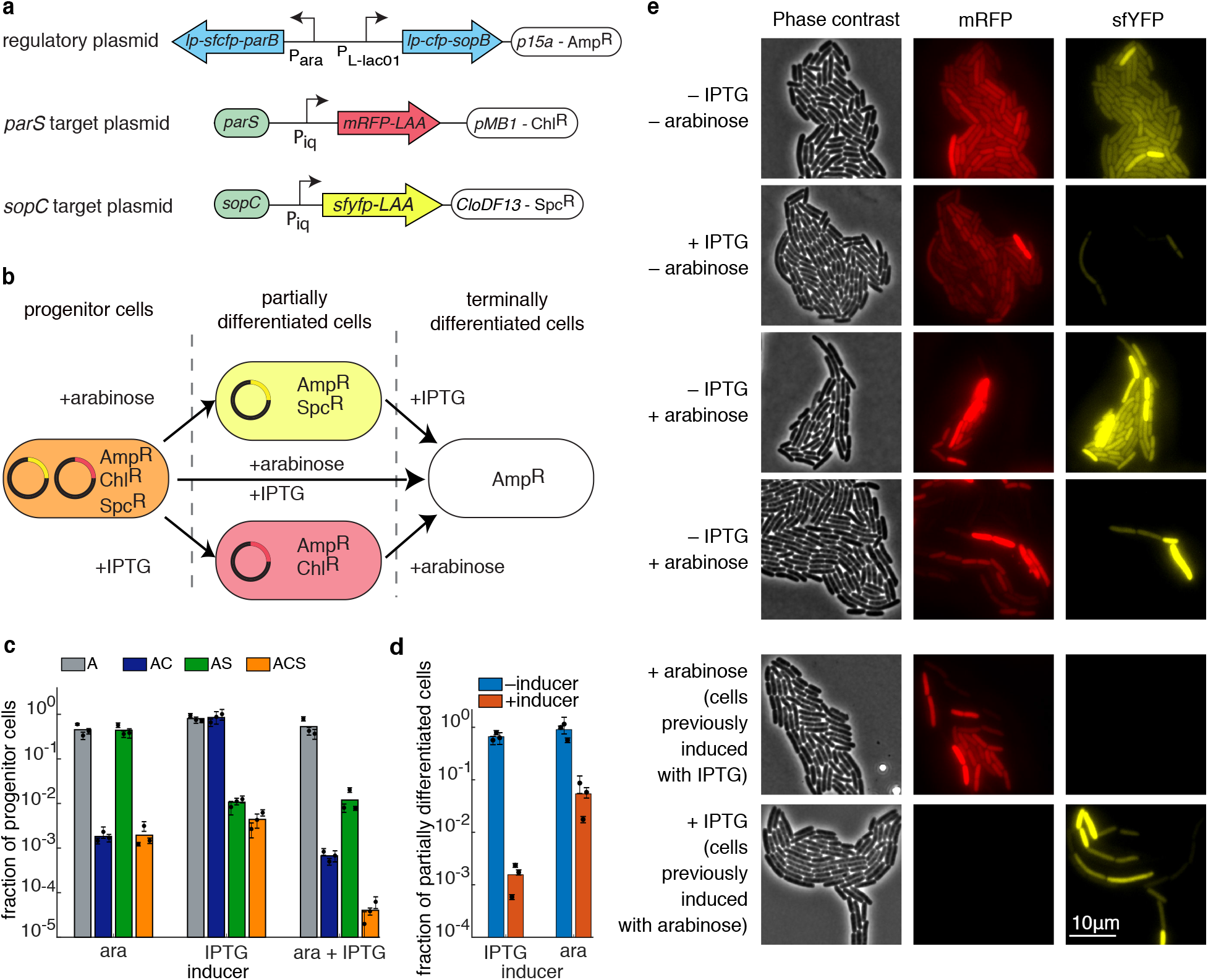
An orthogonal APP system. **(a)** Schematics of key elements on each of the three plasmids used to test orthogonal partitioning. **(b)** Diagram of possible differentiation routes for the progenitor cells. Depending on the inducers, progenitor cells can lose either one of the target plasmids, or both. **(c)** Fraction of progenitor cells in different antibiotic conditions for different combinations of the inducers, as measured by the plating assay (A – ampicillin; C – chloramphenicol; S – spectinomycin). When induced with just arabinose, chloramphenicol resistance is lost. When induced with just IPTG, spectinomycin resistance is lost. When induced with both arabinose and IPTG, both spectinomycin and chloramphenicol resistances are lost. **(d)** Fraction of partially differentiated cells (as measured by the plating assay) with (red) and without (blue) induction of the remaining APP system. In both cases, the remaining APP system retains its ability to differentiate. Colored bars represent the average of all trials, black dots represent the average of each biological replicate and their error bars the standard deviation of technical triplicates (c,d). **(e)** Phase contrast (left), red fluorescence (middle), and yellow fluorescence (right) images of cells after induction with various combinations of inducers. Also shown (bottom two rows) are colonies previously induced for differentiation of one pathway undergoing induction for the other pathway.

We again used the plating assay to assess the amount of differentiation of the circuit after we induced it with one of the two inducers, or both (Fig. 5c). For this plating assay, the resulting cultures were plated after 7 h onto agar with various selective antibiotics (ampicillin (A), ampicillin and spectinomycin (AS), ampicillin and chloramphenicol (AC), or ampicillin, chloramphenicol, and spectinomycin (ASC)) to select for various combinations of plasmids. In each case, the fraction of progenitor cells matched expectations: *e.g*. when only arabinose was used to induce, most cells grew only on ampicillin or ampicillin plus spectinomycin, as the target plasmid conferring chloramphenicol resistance had been lost in the majority of cells.

The above results can also be seen through fluorescence microscopy (Fig. 5e). In the absence of inducer, cells contain both red and yellow fluorescence (top row). However, if one of the inducers is present, the number of cells with the corresponding fluorescence is drastically reduced (second and third rows). If both inducers are present, the resulting population is primarily empty of either fluorescent protein. Note that in each case, there are still progenitor cells present in the culture, indicative of the asymmetric nature of the differentiation. These results also hold for sequential induction, as displayed in the bottom two rows of Fig. 5e. Sequential induction efficiency has been also quantified by plating assay. We picked colonies from cultures induced by IPTG (plates containing ampicillin and chloramphenicol) and arabinose (plates containing ampicillin and spectinomycin) in the assay described in Fig. 5c. Those cells presumable only lost one of the two target plasmids. We inoculated them overnight and restarted the plating assay inducing the culture with the other inducer (the one not used in the previous round of inoculation). The fraction of progenitor cells at steady state in each case is illustrated in Fig. 5d. We can appreciate that both systems are still functioning and able to produce differentiated cells by further APP of the target plasmid that they retained after the first round of induction. Cultures previously induced with IPTG lose efficiency in the second induction with arabinose. The same phenomenon is not observable when cultures are re-induced with IPTG.

## Discussion

In this work, we developed a novel synthetic gene circuit for engineering asymmetric cell differentiation in *E. coli* through the asymmetric partitioning of plasmids. The ParB protein binds a *parS* sequence on the target plasmids, aggregating them into a single nucleoprotein complex and interfering with normal partitioning. It should be noted, however, that the ability of ParB to from the nucleoprotein complex appears to depend on the characteristics of the N-terminal fusion sequence. As we showed, altering the hydrophobicity of the leader peptide or changing the fluorescent protein can alter the system’s ability to perform APP. The fluorescent protein, though, is not a requirement, as a strong homo-oligomerization domain (GCN4zip) is sufficient by itself.

Our circuit distinguishes itself from other synthetic differentiation mechanisms, especially toggle switches, in several important ways. First, differentiation in our circuit occurs through asymmetric cell division, meaning that a progenitor cell will always remain in the culture, ready to reseed the population. Second, differentiation with the APP circuit is irreversible. Once a plasmid is lost in a daughter cell, it cannot be recovered (barring some form of horizontal gene transfer). This means that no refractory period exists if the circuit was used as a memory device. Once a transient signal has been sent, differentiated cells will appear and begin to proliferate as the environment allows. In contrast, when input signals of a toggle switch are transient, the system will reset to its original state after some time^17^. The only way to reset the APP system is to rid the colony of differentiated cells (by whichever means appropriate) and regrow the progenitor cells.

Finally, one disadvantage of corepressive toggle switches is that they are difficult to tune because they generally have a limited parameter space in which they exhibit bistability. The iterative nature of constructing such circuits can add a significant amount of time to the design-build-test cycle^48^. The APP system, though, requires very little tuning, as differentiation requires only the accumulation of the DNA binding protein and not the repression of another transcriptional state. Hence, the balance of two nonlinear processes is unnecessary. This, and the other advantages noted above, make the APP system a great option for creating differentiated multicellular systems from simple prokaryotic hosts.

## Acknowledgements

We are especially grateful to Lucy Shapiro and Christine Jacobs-Wagner for providing plasmids and to Karen Fahrner from the Berg Lab for her help in supplying and consulting on strain HCB84. This work was funded by the Defense Advanced Research Projects Agency (DARPA) Biological Technologies (BTO) Biological Controls Program, award #HR0011-17-2-0012 (approved for public release; distribution is unlimited) (MRB and OI); the National Science Foundation through the joint NSF-National Institute of General Medical Sciences Mathematical Biology Program grant DMS-166290 (MRB); the Welch Foundation grant C-1729 (MRB); and the National Institutes of Health grant R01GM117138 (MRB).

## Author Contributions

MRB, OAI and SM conceived of the study. SM, DLS, and SPB performed experiments and SM analyzed the data. MRB, OAI, and JC oversaw the project. All authors wrote the manuscript.

## Methods

### Plasmids and *E. coli* strains

Plasmids were constructed using Golden Gate assembly, Gibson assembly, and site directed mutagenesis. See Table S2 for the list of plasmids used in this paper. All experiments were conducted in strain JS006ALT (parent strain: *E. coli* K-12 BW25113 Δ*lacI* Δ*araC* (*tetR*^−^) with constitutive *lacI, araC*, and *tetR* cassettes genomically integrated^4,49^), except for the motility assay which required an *E. coli* strain able to swim in soft agarose and was therefore performed in PL64 (parent strain HCB84, *motA* mutant)^46^.

### Plating assay – growth curve

Cells of *E. coli* strain JS006-ALT containing both the regulatory plasmid and the target plasmid were cultured overnight at 37°C shaking at 250 rpm in LB medium containing both ampicillin (Amp, 100 mg/L) and chloramphenicol (Chl, 34 mg/L) antibiotic. Two 250 mL Erlenmeyer flasks containing 50 mL of LB medium with only Amp were inoculated with overnight culture at 0.1% v/v and incubated at 37°C shaking at 250 rpm for up to 8 hours. At 1 hour after inoculation, inducer was added to one of the two cultures – this is what we define as ‘time 0’.

Starting from time 0, the growing culture was sampled every hour. At each sampling, culture was diluted into non-selective LB media and then plated on to two sets of LB agar plates: One set of plates contained only Amp, to assay the total CFU count, and the other set contained both Amp+Chl, to assay the CFU count associated with cells containing target plasmid. 100 *μ*L of each final culture dilution was applied and spread on to each plate using 12 glass beads (Millipore Sigma).

For each time point, culture from each flask was diluted at specific dilution ratio that depended on the presence or absence of inducer and on what antibiotic was present in the LB-agar. For cultures with no added inducer that were plated on solid media plated with or without Chl, and for cultures with added inducer that were plated on solid media without Chl, the dilution ratios are as follows: time 0h: 1/1000; time 1h: 1/2000; time 2h: 1/10000; time 3h: 1/100000; time 4h: 1/200000; time 5-6-7h: 1/500000. For cultures with added inducer that were plated on solid media containing Chl, the dilution ratios are as follows: time 0: 1/1000; time 1h: 1/2000; time 2-3-4-5-6-7h: 1/10000.

For each plate, 12 glass beads were applied to each plate to spread 100 *μ*L of final diluted cell culture. Plates were then incubated at 37°C until colonies reached at least 0.5mm in diameter. Plates were then imaged through Alpha Innotech MultiImage™ light cabinet using AlphaView Fluorchem FC3 (version 3.4.1). Total CFUs on each plate were counted with ImageJ (version 2.0.0-rc-54/1.51g).

All plating assays follow the protocol described above. If only one time point is shown, it is the 7 h post induction time point.

### APP titration assay

Cells were inoculated from overnight culture as described in ‘plating assay’ into four different flasks. At 1 h post inoculation (time 0 in Figure 3) each flask was added with the stated amount of L-arabinose (Millipore Sigma): 0%, 0.002%, 0.02% and 0.2% m/V. 7 hours after induction, cultures were diluted and plated as described in ‘plating assay’. Dilutions for the ParB/parS system: For cultures with no added inducer that were plated on solid media plated with or without Chl, and for cultures with added inducer that were plated on solid media without Chl, the dilution factor was 1 /500000. For cultures with added inducer that were plated on solid media containing Chl, the dilution factor was 1/100000 for cultures induced with 0.002% arabinose, 1/100000 for cultures induced with 0.02% arabinose, 1/10000 for cultures induced with 0.2% arabinose. Dilutions for the SopB/sopC system: For cultures with no added inducer that were plated on solid media plated with or without Chl, and for cultures with added inducer that were plated on solid media without Chl, the dilution factor was 1/500000. For cultures with added inducer that were plated on solid media containing Chl, the dilution factor was 1/100000 for cultures induced with 0.002% arabinose, 1/10000 for cultures induced with 0.02% arabinose, and 1/1000 for cultures induced with 0.2% arabinose.

### APP orthogonal system plating assay

Cells were inoculated from overnight culture as described in ‘plating assay’ into four different flasks. At 1hr post inoculation (time 0 in Figure 3), each flask was added with either no inducer, 0.2% arabinose, 0.1mM IPTG or 0.2% arabinose and 0.1 mM IPTG. 7 hours after addition of inducer, cultures were diluted and plated on to four different sets of plates: Amp, Amp+Chl, Amp and spectinomycin (Spc, 50 mg/L), or Amp+Chl+Spc. The culture with no added inducer was diluted 1/500000 for each plating condition. The culture with 0.2% arabinose was diluted 1/500000 when plated on Amp or Amp+Spc and was diluted 1 /10000 when plated on Amp, Amp+Chl, or Amp+Chl+Spc. The culture induced with 0.1mM IPTG was diluted 1/500000 when plated on Amp, or Amp+Chl, and diluted 1/1000 when plated on Amp+Spc, or Amp+Chl+Spc. Plates were incubated at 37°C until colonies grew to about 0.5 mm in diameter and were imaged as described in ‘plating assay’.

### Single cell Microscopy Assay

We imaged cells incubated underneath a layer of solid LB-1.5% agarose (1–2 mm). 1 *μ*L of cell culture was placed between a slab LB-agarose and a glass cover slip-bottomed 50 mm petri dish with 30 mm glass diameter (MatTek Corporation). Images were acquired using an inverted fluorescence microscope (Nikon) and imaged every 3 min.

### qPCR

Cells were inoculated from overnight culture as described in ‘plating assay’, into two different flasks. At time 0, one hour after the initial inoculation, one flask was induced with 0.2% L-arabinose. Starting at time 0, We sampled each flask every hour. We extracted the following volumes at each time point: time 0 h: 20 mL; time 1 h: 20 mL; time 2 h: 10 mL; time 3 h: 5 mL; time 4 h: 3 mL; time 5 h, 6 h, 7 h: 1 mL. Cells were pelleted and plasmid DNA was extracted with the QIAprep Spin Miniprep Kit at a final elution volume of 50 *μ*L. For the qPCR, we used 1 *μ*L of the DNA resulting from the miniprep. Forward and reverse primers were added to a final total primer concentration of 0.1 *μ*M in addition to 5 *μ*L of Maxima SYBR Green/ROX qPCR Master Mix (Thermo Fisher). Nuclease-free water was added for a final reaction volume of 10 *μ*L. A Bio-Rad CFX96 qPCR machine was used for data collection using the following PCR program: 50°C 2 min, 95°C 10 min, followed by 30 cycles of 95°C 15 s and 60°C 1 min. All of the measurements were followed by melting curve analysis. Results were analyzed using Bio-Rad qPCR analysis software by a relative standard curve. For quantification, a 5-point standard curve covering a 10^4^ fold range of concentrations of the target plasmid and the regulatory plasmid was run in parallel and used to determine the relative DNA abundance in each sample and the efficiency of each primer. It was shown that the qPCR primers for both the target and regulatory plasmid had a primer efficiency between 82.73–93.22%. All of the DNA samples were measured in triplicate, and non-template controls run in parallel to control for contamination and nonspecific amplification or primer dimers. Melting curve analysis was performed to confirm that only a single product was amplified.

The primers used for qPCR in this study were:

Target Plasmid FWD: 5’-GCCGGAAATCGTCGTGGTATTC-3’

Target Plasmid REV: 5’-CAATGAAAGACGGTGAGCTGGTG-3’

Regulatory Plasmid FWD: 5’-CGTCGTTTGGTATGGCTTCATTCAG-3’

Regulatory Plasmid REV: 5’-CTAACCGCTTTTTTGCACAACATGG-3’

### qPCR on individual colonies (colonies)

We collected cells from the induced culture plated without Chl, 7 h after adding L-arabinose as described in ‘plating assay’. Single colonies of the plate were picked and resuspended in 10 *μ*L PBS. Following the same protocol as described in above, 1 *μ*L of the cell resuspension was used for each reaction.

### Colony growth on semi-solid agarose

We inoculated overnight cultures at 30°C of cells transformed with the APP circuit (Fig. 4a) into LB 0.3% m/V agarose plates containing Kan and Spc and 2 mM IPTG in inducing plates. We inserted a pipette tip containing 0.5 *μ*L of overnight culture midway in depth in the LB agarose layer, approximately ~2mm below the surface of the medium, avoiding touching the bottom of the plate. Cells were ejected as the tip was pulled up through the agarose. Plates were incubated at 30°C inverted. Images were obtained after 24, 48 and 72 hours.

### Data Availability Statement

The datasets generated and analyzed during the current study are available from the corresponding author on reasonable request. All plasmids generated during this study are available on AddGene, deposit number 76881.

## Notes

#### Summary of Updates

New updated version of the manuscript.

