## Supplementary Information for "A synthetic system for asymmetric cell division in *Escherichia coli*"

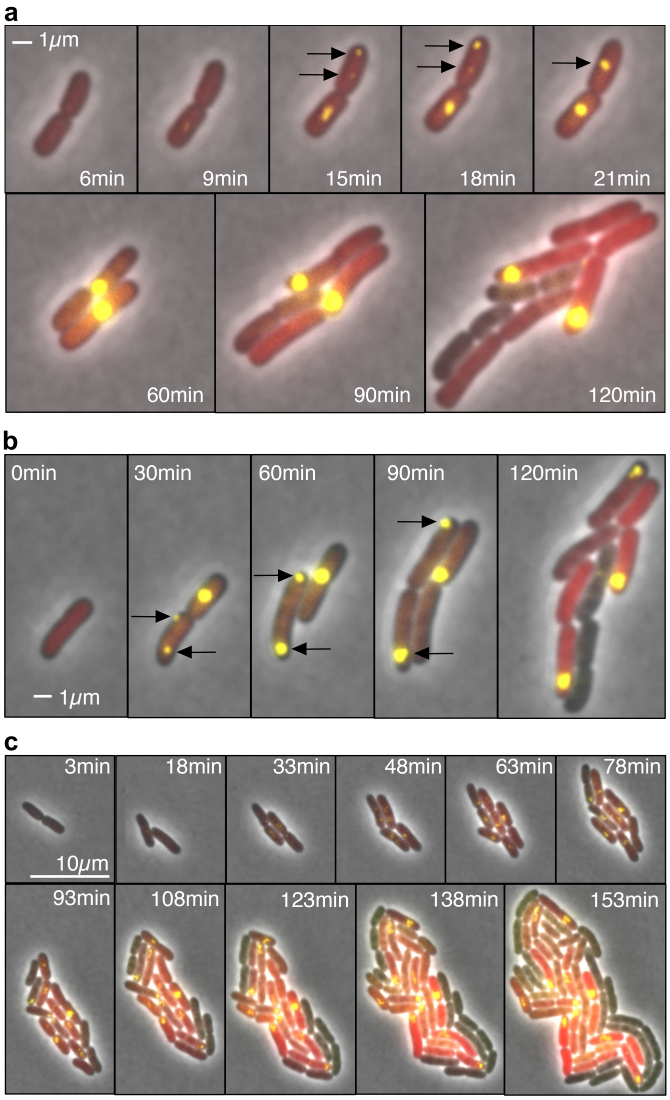


**Supplementary Figure 1.** Time lapse fluorescence microscopy of APP. **(a)** Fluorescence microscopy of cells growing in an agarose pad containing 0.2% arabinose. At early stages of ParB induction, the amount of protein in the cell may not be enough to gather all the plasmids in one cluster, resulting in initial symmetric cell division. Here the two foci indicated by the arrows eventually merge into a single cluster. **(b)** In this example, one cell contains two foci (black arrows). Instead of converging into a single focus, however, the two foci reside at opposite poles of the cell and eventual create two new progenitor cells, each with a single focus. **(c)** This example shows a more complex and chaotic formation of foci. The asymmetric cell division only starts near t=63 min, once each progenitor cell has a single focus. Despite this, approximately 80% of population has differentiated by t=153 min. All the images are acquired under the same conditions.

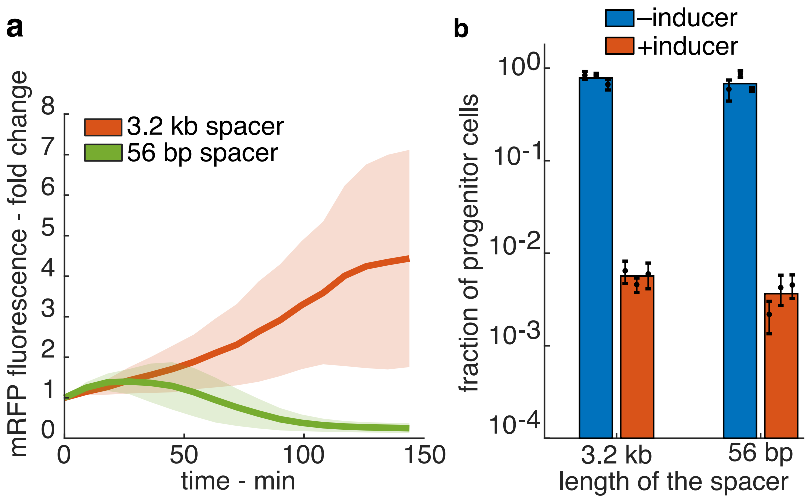


**Supplementary Figure 2.** Spacer length determines ParB dependent gene silencing. **(a)** mRFP fluorescence fold change as a function of time for target plasmids with the *parS* site 3.2 kb (orange) or 56 bp (green) apart from the constitutive promoter driving expression of *mRFP*. Note that red fluorescence is silenced in the absence of the 3.2 kb spacer. Shaded areas represent standard deviation. **(b)** Fraction of progenitor cells in induced and uninduced cultures transformed with an APP system described in (a). Colored bars represent the average of all trials, black dots represent the average of each biological replicate and their error bars the standard deviation of technical triplicates.

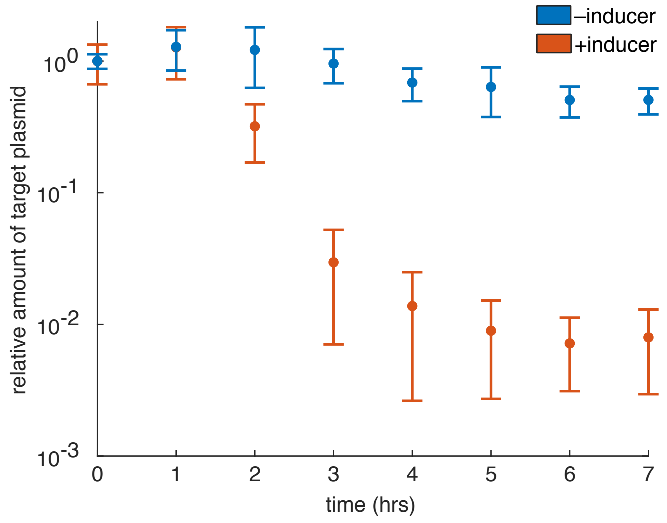


**Supplementary Figure 3.** Relative abundance of the target plasmid compared to the regulatory plasmid as a function of time in both APP induced (orange) and not induced (blue) populations. Error bars represent standard deviation of three biological replicates measured in technical triplicate.

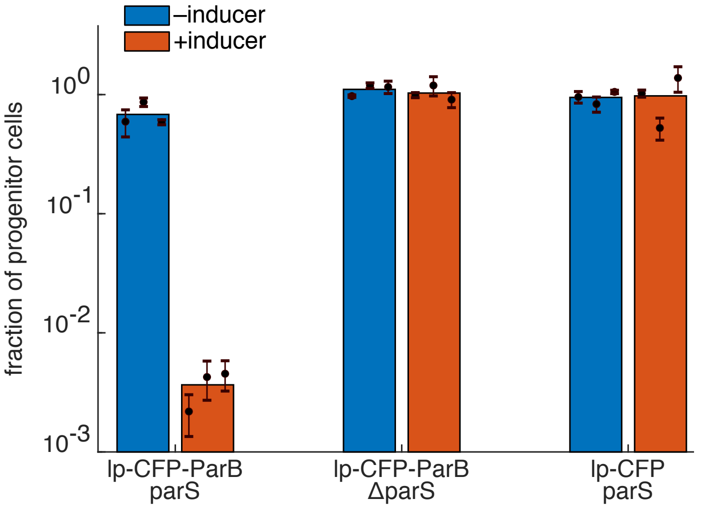


**Supplementary Figure 4.** APP with mutated versions of the system. Fraction of progenitor cells, as measured by plating assay, in induced and not induced cultures. Cells are transformed with an APP system carrying no *parS* on the target plasmid or missing the ParB domain in the regulatory plasmid and compared to the original system (left). This results show that the binding between ParB and *parS* is necessary for APP to happen. Colored bars represent the average of all trials, black dots represent the average of each biological replicate and their error bars the standard deviation of technical triplicates.

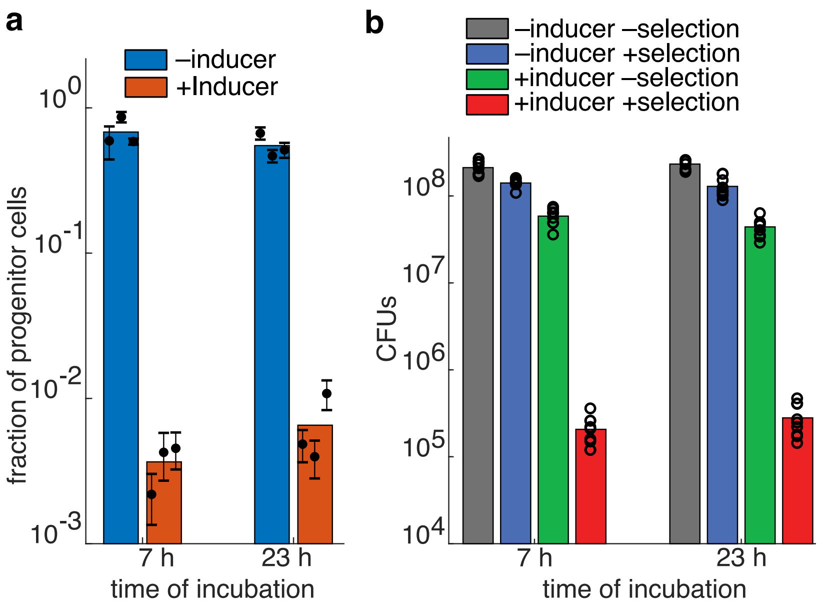


**Supplementary Figure 5.** APP in induced and uninduced population after 24 h from the inoculation. **(a)** Fraction of progenitor cells in induced and uninduced cultures. Colored bars represent the average of all trials, black dots represent the average of each biological replicate and their error bars the standard deviation of technical triplicates. **(b)** Absolute number of CFUs/mL of cultures plated with and without chloramphenicol in induced and uninduced cultures measured by plating assay. Colored bars represent the average of three biological replicates measured in technical triplicates, and black circles are the values of each of those trials.

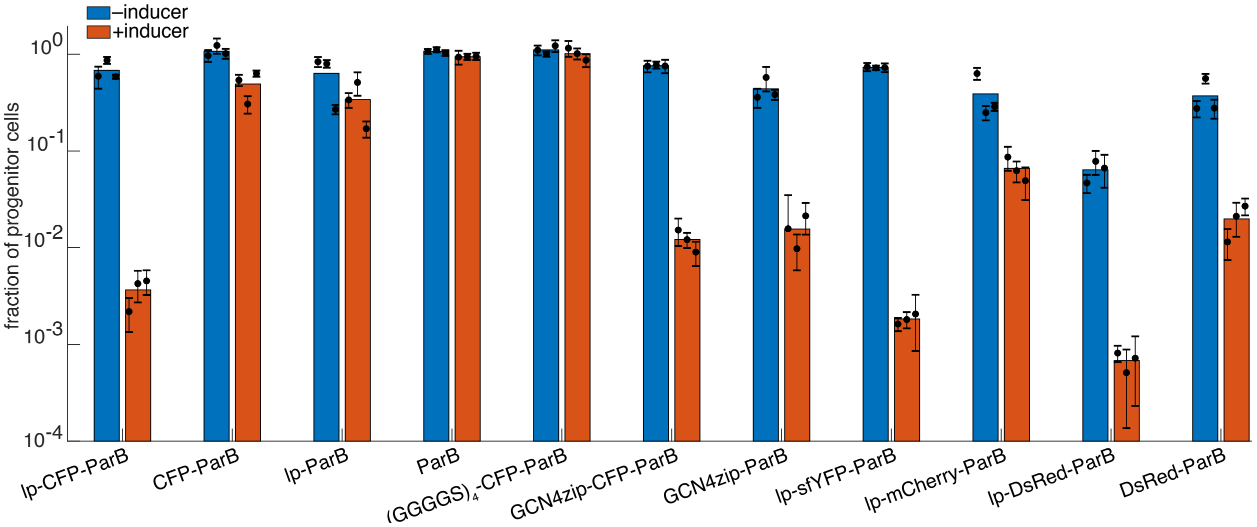


**Supplementary Figure 6.** The fraction of progenitor cells in both induced and uninduced cultures transformed with different versions of the APP systems. In particular, we tested modifications on the leader peptide and fluorescent protein in the ParB construct expressed from the regulatory plasmid. Abbreviations: lp – 17 amino acid leader peptide of the original system, $(GGGGS)_4$ – glycine serine peptide, GCN4zip – GCN4 leucine zipper. Colored bars represent the average of all trials, black dots represent the average of each biological replicate and their error bars the standard deviation of technical triplicates.

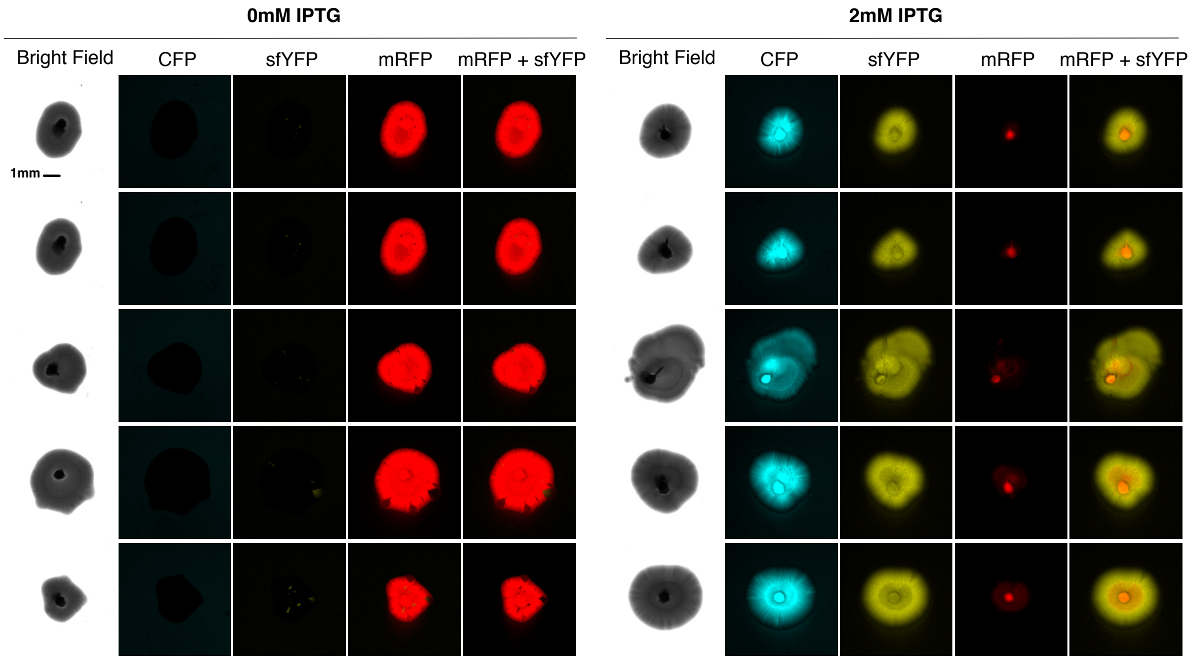


**Supplementary Figure 7.** Pictures of colonies growing in semi-solid agarose plates (0.3%) with (left panel) and without (right panel) inducer. Rows show different colonies all imaged 48 h after inoculation, while columns show different fluorescence channels of the same image. Scale bar is shown only on the picture of the first colony (the bright field) and is the same for every image of the picture.

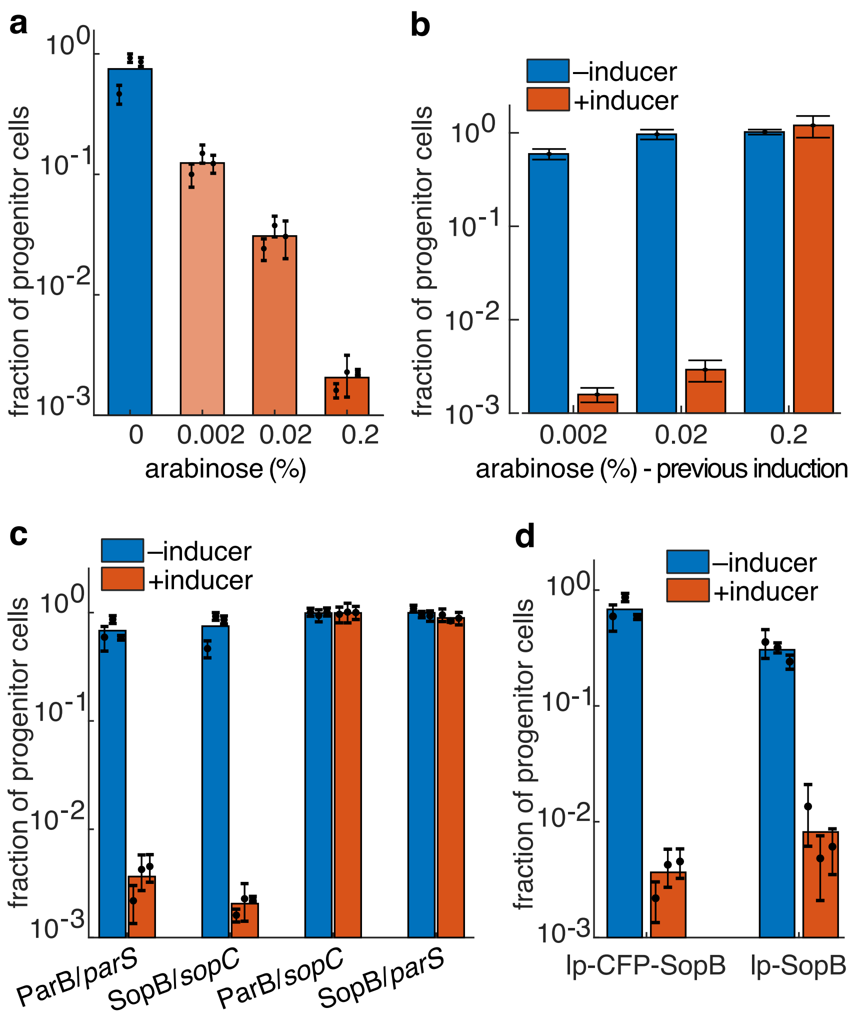


**Supplementary Figure 8.** An orthogonal system for APP. **(a)** Fraction of progenitor cells as a function of arabinose using the SopB/*sopC* system as measured by the plating assay. **(b)** Fraction of progenitor cells in induced (red) and uninduced (blue) culture during the second round of induction (these cells had previously undergone one round of differentiation). Note that a second round of differentiation only occurred if the amount of arabinose used was low. Error bars represent the standard deviation of technical triplicates. **(c)** Fraction of progenitor cells for induced (red) and uninduced (blue) cultures as measured by the plating assay. Each combination of protein and binding site were tested. APP occurred when the protein and binding site came from the same system (left two) and not when they came from different systems (right two). **(d)** Fraction of progenitor cells in induced and uninduced cultures measured by plating assay. We compared systems expressing the original *sopB* construct (*leader peptide-CFP-SopB*) to the one missing the fluorescent tag (leader peptide-SopB). Colored bars represent the average of all trials, black dots represent the average of each biological replicate and their error bars the standard deviation of technical triplicates.

**a**

| ParB/*parS* | Starting Quantity (SQ) ng | | | |
|---|---|---|---|---|
| | regulatory plasmid (Mean) | regulatory plasmid (S.D.) | target plasmid (Mean) | target plasmid (Mean) |
| Colony # 1 | 0.06743 | 0.00295 | 0.00000 | 0.00000 |
| Colony # 2 | 0.11062 | 0.01023 | 0.00000 | 0.00000 |
| Colony # 3 | 0.08388 | 0.00608 | 0.00000 | 0.00000 |
| Colony # 4 | 0.18974 | 0.00693 | 0.00000 | 0.00000 |
| Colony # 5 | 0.48041 | 0.59833 | 0.00000 | 0.00000 |
| Colony # 6 | 0.06343 | 0.00164 | 0.00000 | 0.00000 |
| Colony # 7 | 0.06818 | 0.00040 | 0.00000 | 0.00001 |
| Colony # 8 | 0.09591 | 0.02253 | 0.00000 | 0.00000 |
| Colony # 9 | 0.05233 | 0.00239 | 0.00000 | 0.00000 |
| Colony # 10 | 0.11425 | 0.01053 | 0.00000 | 0.00000 |
| Colony # 11 | 0.04949 | 0.00274 | 0.00000 | 0.00000 |
| Colony # 12 | 0.05474 | 0.00552 | 0.00000 | 0.00000 |

**b**

| SopB/*sopC* | Starting Quantity (SQ) ng | | | |
|---|---|---|---|---|
| | regulatory plasmid (Mean) | regulatory plasmid (S.D.) | target plasmid (Mean) | target plasmid (Mean) |
| Colony # 1 | 0.02470 | 0.00080 | 0.00000 | 0.00000 |
| Colony # 2 | 0.12142 | 0.01055 | 0.00000 | 0.00000 |
| Colony # 3 | 0.15096 | 0.00522 | 0.00000 | 0.00000 |
| Colony # 4 | 0.03575 | 0.00176 | 0.00000 | 0.00000 |
| Colony # 5 | 0.12421 | 0.03078 | 0.00000 | 0.00000 |
| Colony # 6 | 0.05106 | 0.00319 | 0.00000 | 0.00000 |
| Colony # 7 | 0.13096 | 0.01314 | 0.00000 | 0.00000 |
| Colony # 8 | 0.18997 | 0.02643 | 0.00000 | 0.00000 |
| Colony # 9 | 0.07150 | 0.00208 | 0.00000 | 0.00000 |
| Colony # 10 | 0.03715 | 0.00091 | 0.00000 | 0.00000 |
| Colony # 11 | 0.04196 | 0.00087 | 0.00000 | 0.00000 |
| Colony # 12 | 0.08024 | 0.00179 | 0.00000 | 0.00000 |

**Supplementary Table 1.** Starting Quantities of regulatory and target plasmids in colonies picked from the cultures plated at steady state (7 h after induction) without Chl, as measured by qPCR. Data refers respectively to the **(a)** ParB/*parS* and **(b)** SopB/*sopC* system.

| Plasmid | Genes | Ori | Resistance |
|---|---|---|---|
| pAA3 | *Piq - mRFP, parS* | pMB1+rop | Chl |
| pAA41 | *Piq - mRFP, parS* (3.2 kb spacer between *parS* and *Piq*) | pMB1+rop | Chl |
| pAA6 | *Piq- mRFP, parS* | pUC | Chl |
| pCC2 | *Piq - mRFP, ΔparS* | pMB1+rop | Chl |
| pCC28 | *Piq - mRFP, sopC* | pMB1+rop | Chl |
| pCC48 | *Piq - mRFP, BBa_J23100 - phlF, parS* | pMB1+rop | Chl |
| pDD33 | *Para - lp-cfp-sopB* | p15A | Amp |
| pEE29 | *Piq - sfyfp, sopC* | CloDF13 | Spc |
| pEE49 | *Para - lp-sfcfp-parB, Piq - lp-cfp-sopB* | p15A | Amp |
| pSM015 | *Para - lp-cfp-parB** | p15A | Amp |
| pSM018 | *Piq - mRFP, parS* (3.2 kb spacer between *parS* and $Chl^R$) | pMB1+rop | Chl |
| pSM019 | *Para - lp-sfyfp-parB* | p15A | Amp |
| pSM034 | *Para - lp-sopB* | p15A | Amp |
| pSM038 | *PphlF - sfyfp-P2A-motA* | CloDF13 | Spc |
| pSM040 | *Piq - mRFP, parS* | pSC101 | Chl |
| pSM105 | *Para - lp-cfp* | p15A | Amp |
| pSPB845 | *PLlacO1 - lp-cfp-parB, BBa_J23110 - lacI* | p15A | Kan |
| pSPB847 | *Para - parB* | p15A | Amp |
| pSPB849 | *Para - GlySer-cfp-parB* | p15A | Amp |
| pSPB848 | *Para - lp-parB* | p15A | Amp |
| pSPB858 | *Para - GCN4zip.MSE-parB* | p15A | Amp |
| pSPB861 | *Para - lp-mCherry-parB* | p15A | Amp |
| pSPB862 | *Para - DsRed-parB* | p15A | Amp |
| pSPB863 | *Para - lp-DsRed-parB* | p15A | Amp |
| pSPB866 | *Para - GCN4zip.MSE-cfp-parB* | p15A | Amp |
| pSPB867 | *Para - cfp-parB* | p15A | Amp |

**Supplementary Table 2.** Plasmids used in this study. **Ori**: origin of replication. **cfp-parB* genetic cassette has been cloned from the plasmid pNDM220CFPParB[1].
